# Emergent glutamate & dopamine dysfunction in VPS35_(D620N)_ knock-in mice and rapid reversal by LRRK2 inhibition

**DOI:** 10.1101/2024.09.30.615858

**Authors:** A. Kamesh, C.A. Kadgien, N. Kuhlmann, S. Coady, E.P. Hurley, J.C. Barron, M.P. Parsons, A.J. Milnerwood

## Abstract

The D620N variant in Vacuolar Protein Sorting 35 (VPS35) causes autosomal-dominant, late-onset Parkinson’s disease. VPS35 is a core subunit of the retromer complex that canonically recycles transmembrane cargo from sorting endosomes. Although retromer cargoes include many synaptic proteins, VPS35’s neuronal functions are poorly understood. To investigate the consequences of the Parkinson’s mutation, striatal neurotransmission was assessed in 1-, 3- & 6-month-old VPS35 D620N knock-in (VKI) mice. Spontaneous and optogenetically-evoked corticostriatal glutamate transmission was increased in VKI striatal spiny projection neurons by 6 months, when total striatal glutamate release, quantified by iGluSnFR imaging, showed similarities to wild-type. dLight imaging revealed robust increases in VKI striatal dopamine release by 6 months, which were reversed with acute *ex vivo* leucine-rich repeat kinase 2 (LRRK2) inhibition. We conclude that increased glutamate and dopamine transmission in VKI mice progressively emerges in young-adulthood, and that dopamine dysfunction is likely the result of sustained, rapidly-reversible, LRRK2 hyperactivity.

## Introduction

A simplified definition of Parkinson’s disease (PD) is that of a movement disorder, resulting from the death of dopamine neurons in the substantia nigra pars compacta (SNpc). Within ∼4 years of clinical presentation, there is an almost complete loss of nigrostriatal dopamine axon markers. Whereas, the loss of nigral neurons is less severe (30-60%) and remains fairly stable thereafter^1^. The resilience of a population of nigral neurons^1^, and the restorative effect of dopamine replacement therapy (e.g., L-DOPA) against motor symptoms^2^ offer hope that at least some function can be restored in remaining neurons after disease onset.

Unfortunately, no available PD treatment has been shown to slow or prevent disease progression. Furthermore, several disturbances that precede motor dysfunction by many years, e.g., rapid eye movement (REM) sleep behaviour disorder, anosmia, constipation, mood changes, and cognitive decline; are unresponsive to L-DOPA^3–5^. Many neuronal populations, in addition to nigral dopamine neurons, degenerate in PD; notably in the pedunculopontine nucleus and locus coeruleus containing cholinergic and noradrenergic neurons, as well as glutamatergic nuclei in the cortex and thalamus^6–8^. Thus, the evidence implicates multiple neurotransmitter systems in both early symptom manifestation and disease progression.

A point of convergence of multiple neurotransmitter systems in PD pathology are the spiny projection neurons (SPNs) of the dorsolateral striatum^9–11^. SPNs integrate extensive cortical and thalamic glutamate input with modulation by nigrostriatal dopamine to direct behavioural action selection, via downstream basal ganglia nuclei^12^. Too much striatal glutamate and dopamine transmission, however, may be toxic^13–19^ and could act as the initial pathophysiological stress that causes retrograde degeneration and selective loss of SNpc dopaminergic inputs^8,20^.

PD is thought to emerge from the combination of genetic predisposition and environmental stress, especially age^21^. To investigate how these factors precipitate a Parkinsonian state, we developed VKI mice expressing the VPS35 D620N variant linked to clinically-typical PD^22–26^. Heterozygote VKI mice (which model the autosomal dominant presentation in humans) develop tau pathology and nigral loss at >16 months^27,28^. Thus, VKI mice model human PD pathology at advanced ages and are appropriate for the study of early pathophysiological mechanisms.

We previously reported increased glutamate transmission in VKI cortical cultures and increased dopamine release in brain slices of young VKI mice^22,23^. These changes correlated with hyperphosphorylation of leucine-rich repeat kinase 2 (LRRK2) substrates, the kinase believed to be hyperactive in LRRK2-familial and idiopathic PD^21–23,29^. In cultures, despite reversal of LRRK2 substrate hyperphosphorylation, glutamate alterations were resistant to LRRK2 inhibition^23^. Contrastingly, 1-week *in vivo* LRRK2 inhibition rescued decreased dopamine transporter (DAT) protein levels, and normalized dopamine release in VKI brain slices^24^.

Here we sought to define the emergence of glutamate and dopamine synaptic dysfunction as young VKI mice mature, and test whether acute LRRK2 kinase inhibition modifies dopamine release. We found that glutamate and dopamine transmission become elevated by 6 months of age. Elevated dopamine release was rapidly reversed by acute (≥1.5h) LRRK2 kinase inhibition of slices, without evidence of changes to DAT function. We conclude that striatal glutamate and dopamine transmission is progressively increased in VKI mice throughout young adulthood, and that augmented dopamine release is a function of LRRK2 kinase hyperactivity. The data support the argument that LRRK2 inhibition is likely beneficial against synaptic dysfunction in PD and provide a model framework in which to test the neuroprotective potential of LRRK2 inhibitors.

## Methods

### Animals

VPS35 D620N knock-in (VKI) were generated as previously described^22^ and maintained on a C57Bl6/J wild-type (WT) background. Mice were housed and bred in accordance with the Canadian Council on Animal Care regulations (Animal Use Protocol 2017-7888B). All procedures were approved by and governed in accordance with the Neuro Centre of Neurological Disease Models (Animal Use Protocol 2017-7888B) and Memorial University Animal Care Committee (Animal Use Protocol 18-01-MP). 1-, 3-, and 6-month-old male heterozygous VKI and WT mice were used for all measures of spontaneous glutamate transmission. 3- and 6- month male heterozygous VKI and WT mice were used for all experiments requiring stereotaxic injections as these surgeries were performed >4 weeks in advance of experiments.

### Genotyping

Genotyping of all mice was conducted on tail samples at weaning and on post-mortem ear tissue. Samples were digested in 100µL 10% Chelex (Bio-Rad 142–1253; 20 minutes 95°C, 2x vortex) and centrifuged (2 minutes 12,000 RPM) before DNA-containing supernatant (2µL) was added to PCR master mix (18µL, Qiagen 201203: taq polymerase, DNAse and RNAse- free water, 10X buffer, 10mM dNTPs, and custom DNA oligo primers [ThermoFisher: Forward-TGGTAGTCACATTGCCTCTG, Reverse-ATGAACCAACCATCAATAGGAACAC]). DNA was amplified by PCR (program cycle available upon request) and combined with fluorescent DNA intercalating dye (ZmTech LB-001G) prior to iontophoresis. 10µL of PCR product was run in 4% agarose gel and visualized (BioRad UV gel imager) for the presence of 1 or 2 bands to determine WT vs VKI genotype, respectively.

### Surgery

Channelrhodopsin-2 (ChR2), intensity-based glutamate-sensing fluorescent reporters (iGluSnFR), and D1 dopamine receptor-based fluorescent reporters (dLight1.3b) were delivered by stereotaxic injection of AAV constructs as detailed below. 4-6 weeks prior to slice preparation, mice were subcutaneously injected with carprofen (2-4 mg/mL, 20mg/kg, 0.9% NaCl), left to rest for ≥15 minutes, then anaesthetized with isoflurane (5% induction, 1- 2% maintenance) and secured in a stereotaxic head frame (Kopf Instruments). The local analgesic, Marcaine, was injected subcutaneously below the site of incision and hair was removed chemically (Nair) or mechanically (Wahl clippers). An incision was made and skull, leveled using Bregma and Lambda as points of reference. A 0.5mm craniotomy was then created (micro-burr dentistry drill) over the site of injection. A 10µL syringe (Nanofil), attached to a microinjector (Harvard Apparatus Pump11 Elite) was lowered, with coordinates zeroed to the brain surface. Following AAV injection (details below), a 5-minute settling period was allowed before removing the needle. The scalp was then rehydrated with Marcaine, sutured (4-0 silk; Ethicon 683G) and reinforced (Vetbond 3M 1469SB), prior to subcutaneous 0.9% NaCl injection to replace fluids (0.2-0.5mL/10mg). Mice were monitored for pain and discomfort after regaining consciousness and returned to home cage in ∼1 hour. Post-operative monitoring (3 days) included daily subcutaneous carprofen injection.

### Construct-specific injection details

For ChR2 optogenetic stimulation, AAV9-CAG-ChR2-mCherry (Neurophotonics Centre, Université Laval, lot #834 = 4x10^12^ GC/mL) was injected unilaterally (650nL, 1nL/sec) into primary motor cortex (1.5 mm anterior, 1.0 mm lateral, 0.8 mm ventral to Bregma). For optogenetic recording of glutamate release, AAV1.hSyn.iGluSnFr.WPRE.SV40 (Addgene 98929-AAV1, 2.8x10^13^ GC/mL) was injected bilaterally (1uL, 2nL/sec) into the dorsolateral striatum (coordinates: 2.0 mm anterior, 1.0 lateral, 3.2 ventral to Bregma). For optogenetic recording of dopamine release, AAV5-CAG-dLight1.3b (Addgene 125560-AAV5, 1.5x10^13^ GC/mL) was injected bilaterally (300nL, 1nL/sec) into the dorsolateral striatum.

### Preparation of acute brain slices

300µm acute coronal brain slices were prepared from restrained mice which were swiftly decapitated. Brains were quickly transferred to ice-cold recovery solution (1 minute; in mM: 93 NMDG, 93 HCl, 2.5 KCl, 1.2 NaH_2_PO_4_, 30 NaHCO_3_, 20 HEPES, 25 glucose, 5 sodium ascorbate, 3 sodium pyruvate, 10 MgSO_4_·7H_2_O, 0.5 CaCl_2_·2H_2_O, pH 7.3-7.4, 290-310 mOsm; carbogen-infused). Following this, extra-striatal rostral and caudal brain regions were removed by coronal sectioning, and the remaining block was mounted onto a vibrating-blade microtome platform with sodium acrylate (Leica Microsystems VT 1200S). After hemisection at the midline, coronal sections were cut, and slices were then transferred to warm recovery solution (35°C, 15 minutes). A final transfer of slices into holding chambers containing room-temperature artificial cerebrospinal fluid (aCSF; in mM: 125 NaCl, 2.5 KCl, 25 NaHCO_3_, 1.25 NaH_2_PO_4_, 2 MgCl_2_, 2 aCl_2_, 10 glucose, pH 7.2-7.4, 300-310 mOsm, 22-25°C, carbogen-infused) was followed by ≥45 minutes settling period before recording in aCSF began.

### Whole-cell patch-clamp electrophysiology

Whole-cell patch-clamp electrophysiology was used to obtain recordings of spontaneous and ChR2-evoked glutamate transmission in SPNs of the dorsolateral striatum (caudate/putamen), as previously^30^. Slices were transferred to a recording chamber perfused with aCSF (22-25°C) containing 100µM picrotoxin (Tocris 1128) at a flow rate of ∼1.5mL/min. Slices were visualized on an Olympus BX51 microscope (40x magnification, 2x digital zoom, IR-DIC and Q-Imaging Electro camera). SPNs were identified within the dorsolateral striatum based on somatic size (8-20µM) and distinct morphology, 50-150µM below the surface of the slice. Borosilicate glass capillary tubes (Harvard Apparatus 640805) were pulled using a Sutter P-1000 micropipette puller to form 1µm tip recording pipettes (filled resistance 4-8 MOhms; in mM: 130 Cs methanesulfonate, 5 CsCl, 4 NaCl, 1 MgCl_2_, 5 EGTA, 10 HEPES, 5 QX-314, 0.5 GTP, 10 Na_2_ phosphocreatine, 5 MgATP, and 0.1 spermine, pH 7.2, 290mOsm). A motorized micromanipulator (Sutter Instrument MP-285) was used to patch onto identified SPNs, with signals obtained by MultiClamp 700B amplifier in voltage-clamp configuration, filtered at 2kHz and digitized at 10kHz (Molecular Devices Axon Digidata 1440A). Membrane properties were determined with the membrane-test function while holding at -70mV. Synaptic recordings were initiated after a 2-minute settling period, with access resistance tolerance set to 27MΩ and recordings discarded if Δ≥10% over the elapsed period. Spontaneous excitatory postsynaptic currents (sEPSCs) were recorded over a minimum 2-minute period at -70mV (gain 20) and analyzed in Clampfit10 (Molecular Devices; 5pA peak event threshold, confirmed manually). Non-unitary events were retained for inter-event interval analysis, but only unitary events were used for amplitude and decay constants. Cumulative distributions were used to evaluate amplitude and inter-event intervals in each recorded SPN. Unitary events in each recording were averaged and decay tau measured by 1-term exponential fit.

ChR2-evoked post-synaptic currents (ChR2-PSCs) were stimulated by wide-field illumination with blue light (473nm, XCite Series 120Q) transmitted through the 40x immersion objective generating 2.7 mW. 5ms pulses were controlled by a Lambda SC Smart Shutter Controller (Sutter Instruments) administered in trains of 4 pulses with 100ms inter-pulse intervals. Trains were repeated every 30 seconds, 5-10 times and averaged for analysis. Recordings of ChR2-PSCs mediated by α-amino-3-hydroxy-5-methyl-4-isoxazolepropionic acid receptors (AMPARs) were taken at -70mV (gain 2) averaged across 5 repetitions. AMPAR and N-methyl-D-aspartate receptor (NMDAR) currents were generated by single 5ms pulse stimulations repeated 5x every 30 seconds at -70mV then +40mV to assess AMPAR+NMDAR-mediated ChR2-PSCs, respectively. Peak NMDAR current was estimated at +40mV, 40ms post AMPA peak, and AMPAR currents were then isolated at +40mV by bath application of 10µM D-APV (NMDAR blocker, Tocris 0106). AMPA rectification indices were calculated as a ratio of isolated AMPAR ChR2-PSC peak at +40mV : -70mV.

### iGluSnFR and dLight imaging

Imaging of iGluSnFR and dLight recordings were obtained using a 2X objective focused on dorsolateral STR. Slices were wide-field illuminated with blue light (473nm, CoolLED pE-340fura), with the shutter, EM-CCD camera (Andor iXon Ultra 897), and stimulus isolator (WPI A365) triggered by Clampex & Digidata 1550B). The stimulus isolator was connected to a monopolar tungsten stimulating electrode (A-M Systems 574000) lowered into dorsolateral striatum 50-100µM beneath the surface of the tissue. Recordings were captured using Andor Solis software using 4x4 binning at 205Hz. For these experiments, spontaneous iGluSnFR & dLight transients were excluded from analysis of responses to electrical stimulation. For iGluSnFR recordings, a stimulation train was delivered at fixed stimulation intensity (150uA train of 10 x 0.2ms pulses with 100ms inter-pulse intervals). No-stimulation (for background & bleach subtraction) and stimulation trials were alternated at 1-minute intervals, repeated every 2 minutes, until 5 no-stimulation trials were recorded. An 11^th^ pulse was delivered with each repetition of the stimulation trial, with increasing inter-pulse intervals between 500-5000ms. The same protocol was used to obtain pulse train dLight recordings after 2-pulse dLight stimulation with 4s inter-pulse intervals, delivered at 50- 400uA, every 2 minutes to generate stimulus response curves. Responses were then assessed to determine the stimulus intensity corresponding to 50-70% maximum response (FIJI software) for subsequent pulse train dLight experiments. iGluSnFR and dLight videos were converted to the change in fluorescence intensity over time (ΔF/F; FIJI software) with peak and decay analyzed in Clampfit (decays measured by the 1-term exponential function of each trace). Absolute and normalized peak and decay of pulse train responses were derived from the average of 4 responses.

### Acute LRRK2 kinase inhibition

Slices from 6-month-old mice were pre-incubated ≥1.5hours and then bath-perfused during dLight recording with LRRK2 kinase inhibitor MLi-2 (500 nM, Tocris 5756, 45% Captisol® in PBS) or vehicle-control (45% Captisol® in PBS 377.6µL/L aCSF). Captisol® vehicle was also present in all 3-month-old dLight recordings to be able to compare dLight recordings across age.

### Statistical Analysis

All data presentation and statistics were conducted in GraphPad Prism10. Data were tested against ROUT outlier analysis with a liberal maximum false discovery rate Q=1%. Rarely, identified outliers were suppressed from the dataset prior to parametric or non-parametric distributions testing (D’Agostino and Pearson). Unpaired t-test/1-way ANOVA (if parametric) or Mann-Whitney U-test/Kruskal-Wallis testing (if non-parametric) were used to analyse data. *Post-hoc* analysis was performed if ANOVA *p*<0.05 by Tukey’s (parametric) or Dunn’s (non-parametric) multiple comparisons tests as appropriate. All 2-way ANOVA comparisons used Šídák post-hoc analyses. Asterisks represent *p*<0.05 from pair-wise comparisons. All statistical analyses are described in figure legends, and *post-hoc* analyses are reported in Tables S1 & S2. Trends are defined as 0.10 > *p* > 0.05. Data are presented as n=observations from (n) animals (e.g., WT = 6(3) is 6 observations from 3 WT animals).

## Results

### VKI SPNs show progressive increase in spontaneous activity by 6 months

We compared spontaneous excitatory postsynaptic currents (sEPSCs) by whole-cell patch-clamp as before^30^ in VKI and WT SPNs from 1-, 3-, and 6-month-old mice (Figure 1 A). Although similar at 1 month, we found that VKI sEPSCs begin to diverge from WT at 3 months with larger amplitudes.

**Figure 1.**
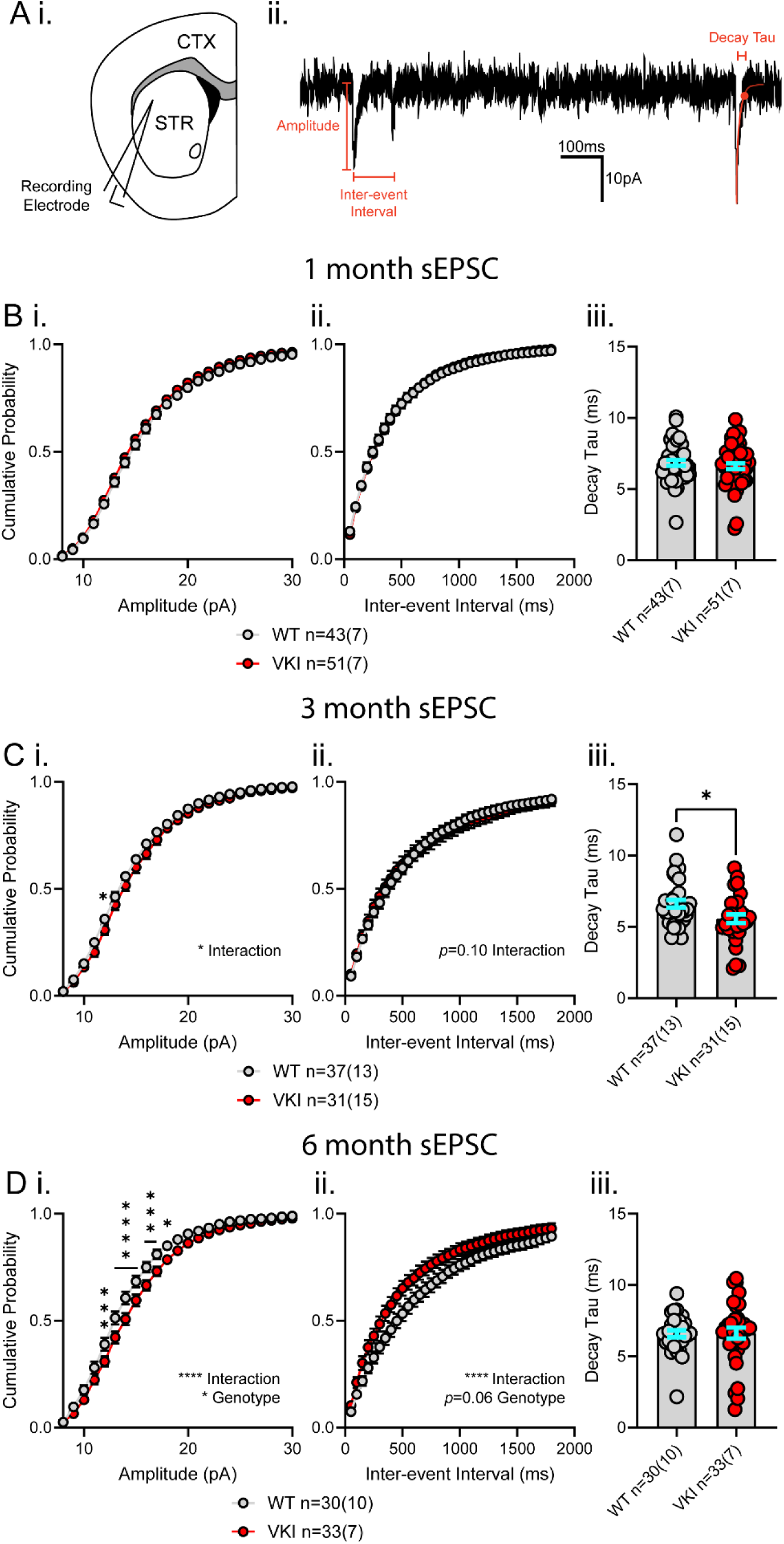
Spontaneous glutamate transmission increase in dorsolateral striatal VKI SPNs emerges by 6 months. **A i)** Schematic depicting whole-cell patch-clamp recording of spiny projection neurons (SPNs) in the dorsolateral striatum (STR) from acute coronal brain sections also containing cortex (CTX). **ii)** Representative trace of voltage-clamp recording of spontaneous excitatory post-synaptic current (sEPSC) from dorsolateral striatal SPNs. Representations of amplitude, inter-event interval, and decay tau measurements annotated in red. **B)** At 1 month, **i)** cumulative probability distributions of sEPSC amplitude are not significantly different between VKI and WT SPNs (2-way ANOVA Interaction *p*>0.99, genotype *p*=0.22). **ii)** Cumulative probability distributions of inter-event intervals are not significantly different between VKI and WT SPNs (2-way ANOVA interaction *p*=0.99, genotype *p*=0.93). **iii)** sEPSC decay tau is not significantly different between VKI and WT SPNs (Unpaired t-test *p*=0.43). **C** At 3 months, **i)** cumulative probability distributions show VKI SPNs have more, larger amplitude sEPSC than WT SPNs (2-way ANOVA interaction *p*=0.02, genotype *p*=0.21). **ii.** sEPSC inter-event intervals are not significantly different between genotypes (2-way ANOVA interaction *p*=0.10, genotype *p*=0.95). **iii.** Average sEPSC decay tau is significantly faster in VKI vs WT neurons (Mann-Whitney U-test *p*=0.01). **D** At 6 months, **i)** cumulative distributions show a significantly higher number of larger-amplitude sEPSCs in VKI neurons (2-way ANOVA interaction *p*<0.0001, genotype *p*=0.02). **ii.** Cumulative distributions of sEPSC inter-event intervals indicate VKI neurons have shorter inter-event intervals than WT neurons (2-way ANOVA interaction *p*<0.0001, genotype *p*=0.06). **iii.** Average sEPSC decay taus are not significantly different in VKI vs WT neurons (Mann-Whitney U-test *p*=0.65). Asterisks denote pairwise comparisons; * = 0.05>*p*>0.01, ** = 0.01>*p*>0.001, *** = 0.001>*p*>0.0001, **** = *p*>0.0001. 2-way ANOVA comparisons listed if *p*<0.10.

VKIs exhibit robustly elevated amplitude and frequency (ie. reduced inter-event interval) by 6 months (Figure 1B-D). The decay of sEPSCs was faster in VKI SPNs at 3 months but normalized to WT levels at 6 months. An age-dependent decrease in WT sEPSC amplitude and frequency was absent in VKI SPNs (Table S2).

We also observed early reductions in membrane tau of 1-month-old VKI SPNs, that normalized to WT levels by 3 months (Figure S1 A,B,C iii). VKI SPNs also trend towards increased membrane capacitance at 3- and 6-months, but not at 1 month (Figure S1 A,B,C i). Membrane resistance was reduced by 6 months in VKI SPNs (Figure S1 A, B, C, ii). As with sEPSC properties, membrane capacitance and resistance measures were altered with age within WT SPNs, but not in VKI neurons (Table S2). Together, age-dependent changes to passive membrane properties, and a progressive (relative) increase in the amplitude, frequency, and decay of spontaneous glutamate transmission is observed in VKI SPNs.

### Corticostriatal glutamate transmission is progressively increased in VKI mice by 6 months

To determine whether increased frequency and amplitude of sEPSCs are due to changes in corticostriatal inputs, optogenetic stimulation of glutamate release onto patched SPNs was conducted at 3 and 6 months (Figure 2 A i). Light stimulation of local cortical terminals generated ChR2-induced post-synaptic currents (ChR2-PSCs). Glutamatergic currents through AMPA and NMDA receptors were isolated by membrane gating properties (ie. holding potential) and by pharmacology with the NMDAR blocker, D-AP5 (Figure 2 A ii). At 3 months, we found no change to VKI AMPAR- or NMDAR-mediated current amplitude. The ratio of NMDA:AMPA current and AMPAR- rectification indices were likewise similar (Figure 2 B). In contrast, we found AMPAR and NMDAR currents were both (equally) elevated in 6-month-old VKIs (Figure 2 C i-ii). There was no change in NMDA:AMPA ratio, or AMPA rectification index at this age (Figure 2 C iii-iv).

**Figure 2.**
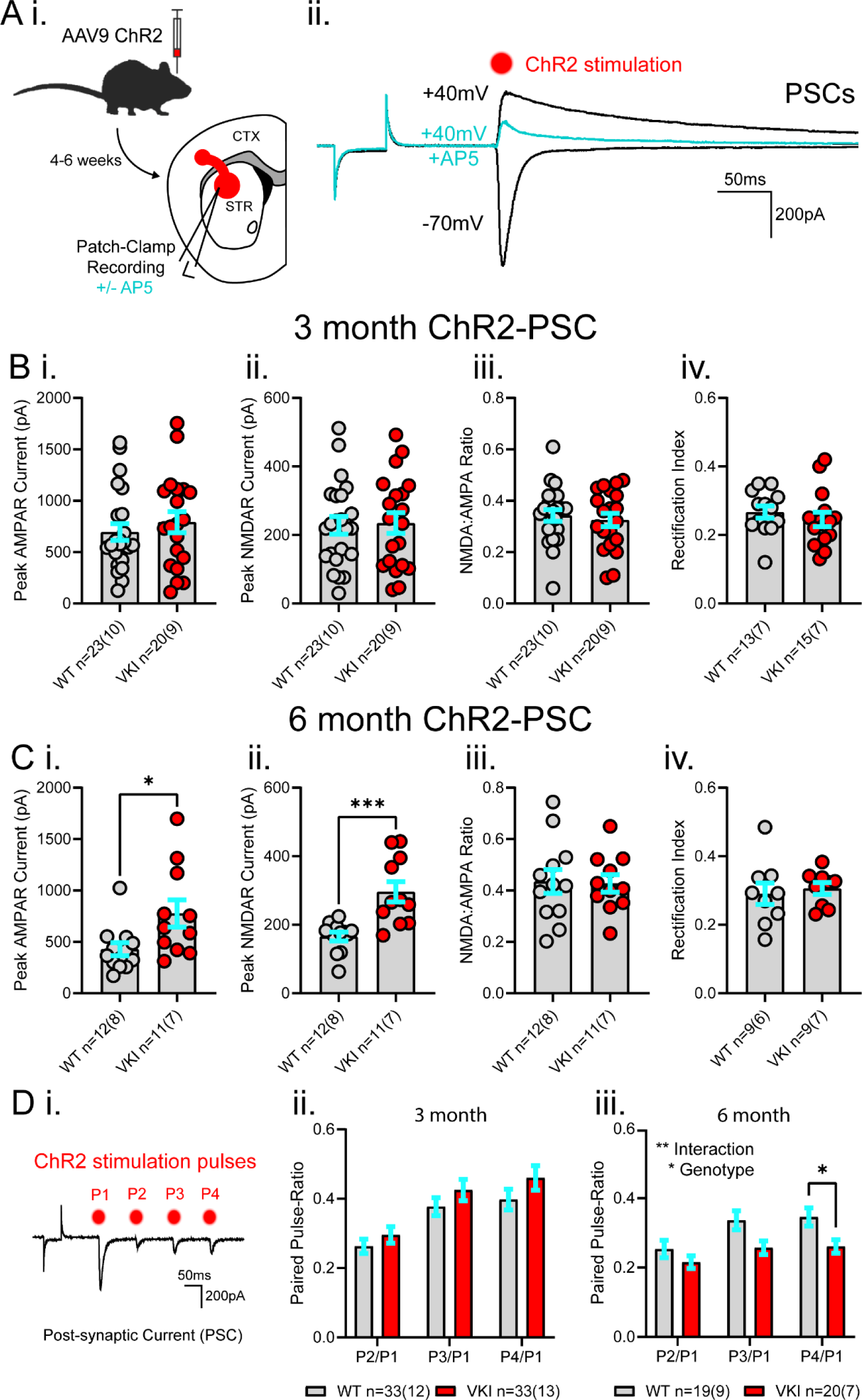
Evoked corticostriatal glutamate transmission increases in VKI SPNs by 6 months. **A i)** Schematic depicting ChR2 viral injection into motor cortex 4-6 weeks prior to preparation of acute coronal slices containing CTX and STR. Whole-cell patch-clamp recordings were obtained from dorsolateral striatal SPNs +/- NMDAR inhibitor, APV. **ii)** Representative ChR2-PSC traces in response to single pulses of light. SPNs that were held at membrane potentials of -70mV or +40mV (in black), and +40mV in the presence of AP5 (in cyan). **B** At 3 months, **i)** the peak amplitude of AMPAR-mediated currents at -70mV are not significantly different between WT and VKI SPNs (Unpaired t-test *p*=0.46). **ii)** Amplitude of NMDAR-mediated currents are not significantly different between genotypes at +40mV (Unpaired t-test *p*=0.87). **iii)** Ratios of +40mV NMDAR current to -70mV AMPAR current are not significantly different between WT and VKI SPNs (Unpaired t-test *p*=0.60). **iv)** Rectification Index of AMPA current is not significantly different between WT and VKI SPNs (Unpaired t-test *p*=0.45). **C** At 6 months, **i)** VKI SPNs show larger currents than WT SPNs, in peak amplitude of both AMPAR-mediated current (Mann-Whitney U-test *p*=0.01) and **ii)** NMDAR-mediated current (Unpaired t-test *p*=0.0005). **iii)** Ratio of NMDA:AMPA currents are not significantly different between genotypes (Unpaired t-test *p=*0.90). **iv)** AMPA Rectification Index is not significantly different between WT and VKI SPNs (Unpaired t-test *p*=0.67). **D i)** Representative ChR2-PSCs with 4 pulses of ChR2 stimulation (in red), 100ms inter-pulse interval. **ii)** At 3 months, the paired-pulse ratio (PPR) of 4 responses to ChR2 stimulation, normalized to the first response, is not significantly different between VKI and WT SPNs (2-way ANOVA interaction *p*=0.34, genotype *p*=0.22). **iii)** At 6 months, the paired-pulse ratio is significantly reduced in VKI vs WT SPNs (2-way ANOVA interaction *p*=0.001, genotype *p*=0.04).

To assess presynaptic probability of release, we quantified paired-pulse ratios (PPRs) at 100ms intervals (Figure 2 D). There was no significant difference to PPRs in VKI neurons at 3 months, but a significant reduction in PPR was observed in 6-month-old VKI SPNs, indicative of elevated probability of presynaptic release (Pr). Taken together, the data show corticostriatal glutamate transmission becomes elevated in VKI SPNs, relative to WT littermates, and that this correlates with an equivalent increase in both AMPAR and NMDAR transmission.

### Total striatal glutamate release appears reduced in 3-month-old VKIs and normalizes to WT levels by 6 months

In addition to changes in presynaptic release, glutamate currents recorded in SPNs are modulated by regulation of postsynaptic responsiveness, thalamic glutamate, midbrain dopamine, and GABAergic interneuron activity^31–38^. To directly assess presynaptic release, extracellular glutamate transients were quantified with the intensity-based glutamate-sensing reporter, iGluSnFR1 (Figure 3 A) in response to fixed intensity pulse-trains (10x10Hz) delivered by local striatal electrical stimulation. The amplitude of iGluSnFR responses was markedly reduced in 3-month-old VKI striata, and there was a significant interaction between the normalized peak and pulse number (Figure 3 B i-ii and Table S1). The recovery capacity of iGluSnFR responses following the pulse train was similar at 3 months, as was the decay of the pulse train response (Figure 3 B iii-iv). At 6 months, iGluSnFR responses were similar to WT responses across all measures (Figure 3 C). Interestingly, a significant age-dependent reduction in the amplitude of iGluSnFR transients was only observed for WT and not VKI responses (Table S2). No age dependent changes were observed in other parameters except for the decay of iGluSnFR responses in VKI striata between 3 and 6 months, which appear significantly increased with age. In summary, total striatal glutamate release is reduced in VKI mice at 3 months, but similar to WT at 6 months, suggesting that total presynaptic glutamate release is relatively increased in the VKI striatum with age.

**Figure 3.**
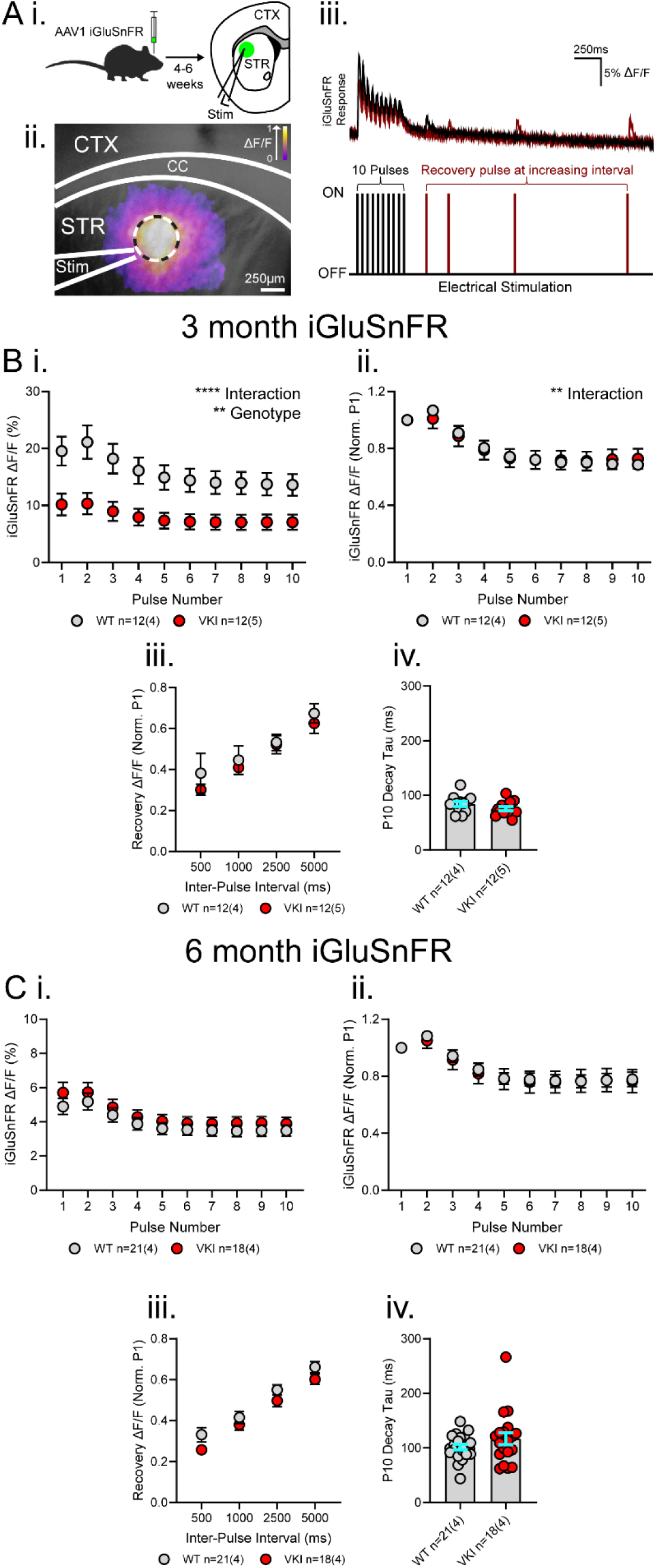
Total striatal glutamate release in VKIs normalize to WT levels by 6 months. **A i)** Graphic depicting viral injection of iGluSnFR in dorsolateral striatum 4-6 weeks prior to acute slice preparation. **ii)** Area of iGluSnFR expression in coronal slices containing CTX and STR depicted in green along with stimulus electrode (Stim) placed in dorsolateral STR. **iii)** Visualization of intensity-based glutamate-sensing fluorescent reporter (iGluSnFR) change in fluorescence relative to baseline (ΔF/F) in dorsolateral STR in response to local electrical stimulation (Stim). Low to high ΔF/F as a ratio of 0 to 1 represented as a gradient from purple to white, respectively. Boundaries of the corpus callosum (CC) separate CTX and STR in a coronal brain slice. Dashed circle represents region of interest (ROI) of 20 pixels used for iGluSnFR analysis. **ii)** Representative iGluSnFR response corresponding to trains of 10x10Hz pulses of electrical stimulation followed by 11^th^ pulse at increasing interval with each of 4 repetitions. Black trace represents 1^st^ of 4 stimulations with subsequent stimulations overlayed in maroon. **B** At 3 months, **i)** VKI vs WT iGluSnFR response size (ΔF/F) is significantly smaller in VKI brain slices (2-way ANOVA interaction *p*<0.0001, genotype *p*=0.007). **ii)** When normalized to 1^st^ pulse (Norm. P1), VKI iGluSnFR responses are significantly different from WT (2-way ANOVA interaction *p*=0.006, genotype *p*>0.99). **iii)** VKI and WT recovery after 10x10Hz stimulation, normalized to 1^st^ pulse, are similar (2-way ANOVA interaction *p*=0.84, genotype *p*=0.50). **iv)** Decay of the last response in pulse train (P10) is similar between VKI and WT iGluSnFR responses (Unpaired t-test *p*=0.17). **C** At 6 months, **i)** VKI vs WT iGluSnFR response size is similar in VKI and WT brain slices (2-way ANOVA interaction *p*=0.90, genotype *p*=0.40). **ii)** Normalized VKI iGluSnFR responses are not significantly different from WT (2-way ANOVA interaction *p*=0.77, genotype *p*=0.87). **iii)** VKI recovery capacity after 10x10Hz stimulation is similar to WT capacity (2-way ANOVA interaction *p*=0.63, genotype *p*=0.12). **iv)** Decay of the last response to 10x10Hz stimulation is not different between VKI and WT iGluSnFR responses (Mann-Whitney U-test *p*=0.33).

### Increased striatal dopamine release appears in 6-month-old VKI mice

We measured extracellular dopamine transients in the dorsolateral striatum using the dLight fluorescence reporter, evoked by electrical stimulation in slices from 3- and 6-month-old mice (Figure 4-6). We did not observe any changes to dLight amplitudes, or responses to pulse-train stimulation in 3-month-old VKI striata (Figure 4 A); however, recovery capacity following pulse train stimulation was significantly reduced in VKIs.

**Figure 4.**
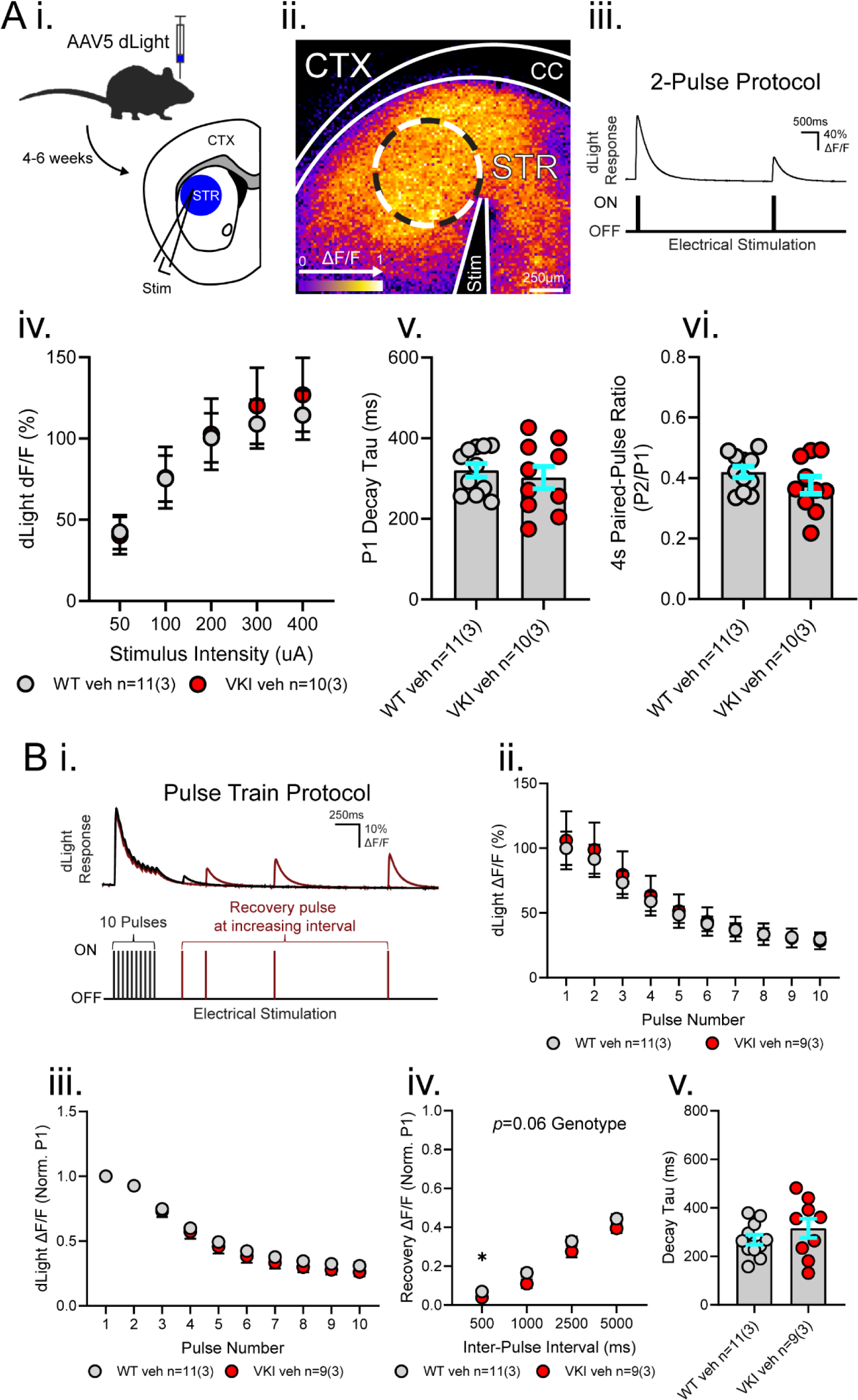
dLight responses to electrical stimulation are not altered in VKI brain slices at 3 months. **A i)** Graphic depicting stereotaxic injection of AAVs encoding for dLight in dorsolateral STR 4-6 weeks prior to acute slice preparation. Area of dLight expression in coronal slices containing CTX and STR depicted in blue with Stim placed in dorsolateral STR. **ii)** Visualization of dLight change in fluorescence from baseline (ΔF/F) in in response to local electrical stimulation (ΔF/F values of 0 to 1 represented as a gradient from purple to white, respectively). Boundaries of the CC separate CTX and STR in a coronal brain slice. Dashed circle represents ROI of 40 pixels used for dLight analysis. **iii)** Representative dLight response in dorsolateral striatum corresponding to 2 pulses delivered with 4s inter-pulse interval. At 3 months **iv)** Increasing stimulus intensity did not alter size of dLight responses in VKI vs WT STR (2-way ANOVA interaction *p*=0.69, genotype *p*=0.83). **v)** Decay of the 1^st^ response peak (P1) is similar between VKI and WT brain slices (Unpaired t-test *p*=0.57). **vi)** PPR at 4s (P2 Peak/P1 Peak) is similar between VKI and WT brain slices (Unpaired t-test *p*=0.21). **B i)** Representative dLight response in dorsolateral STR corresponding to 4x stimulation with 10x10Hz pulse train and subsequent recovery stimulation. At 3 months **ii)** dLight responses were not significantly different in size between VKI and WT brain slices (2-way ANOVA interaction *p*=0.99, genotype *p*=0.85). **iii)** dLight responses relative to 1^st^ pulse (P1) were not different between VKIs and WTs (2-way ANOVA interaction *p*=0.69, genotype *p*=0.45). **iv)** Recovery capacity following 10x10Hz stimulation are lower in VKI brain slices (2-way ANOVA interaction *p*=0.81, genotype *p*=0.06). **v)** Decay of the pulse train response is not significantly different in VKI vs WT brain slices (Unpaired t-test *p*=0.28).

At 6 months, dLight response peaks were significantly increased in VKI brain slices, without any change in decay tau, or PPR at 4s intervals (Figure 5 and Table S1). Pulse train response peaks in VKIs were accordingly increased, without changes in pulse-train response pattern (Figure 6).

**Figure 5.**
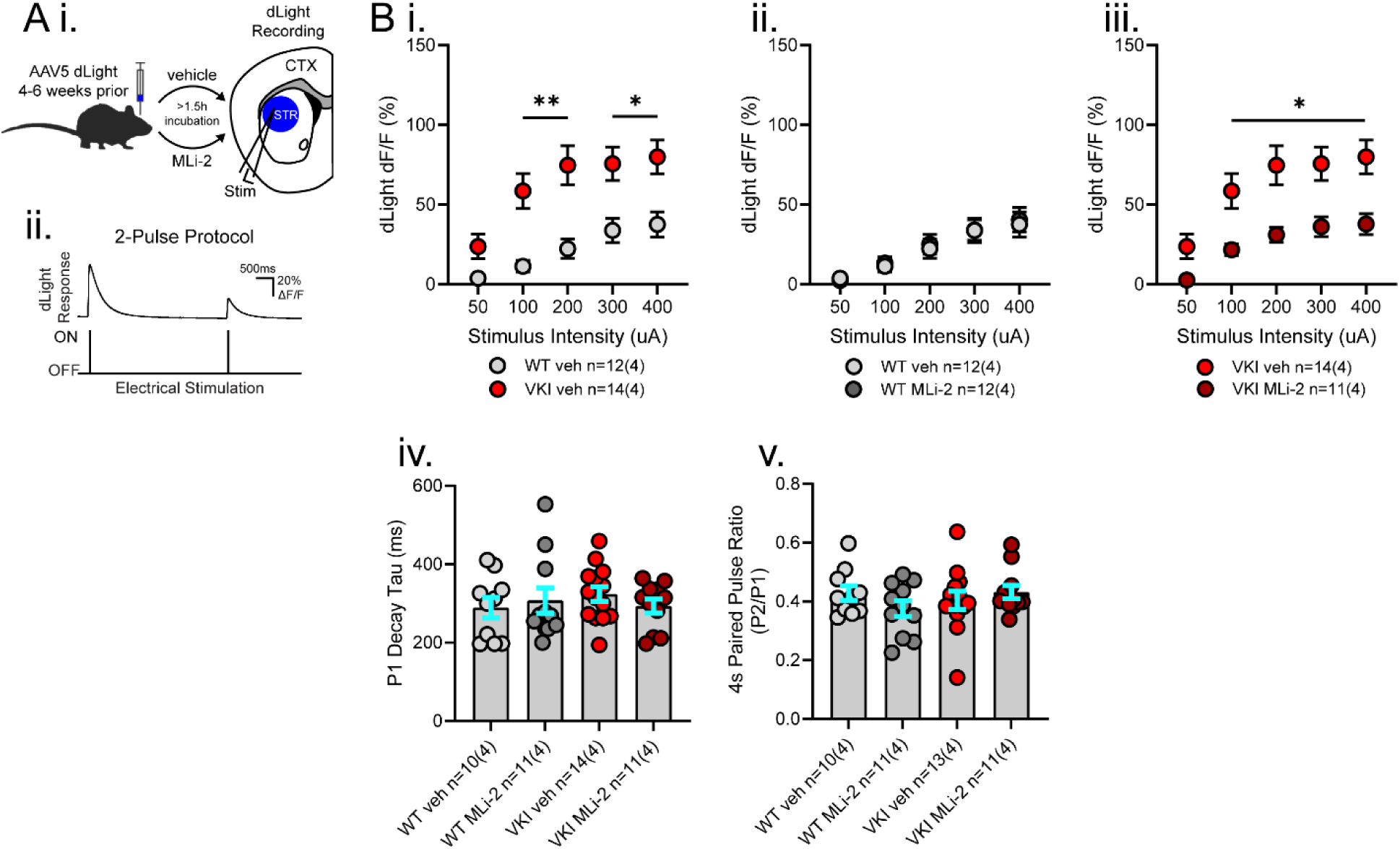
Dopamine release in VKI brain slices at 6 months is elevated and can be reduced with acute MLi- 2 treatment. **A i)** Graphic depicting stereotaxic injection of AAVs encoding for dLight into the dorsolateral striatum 4-6 weeks prior to acute slice preparation. Coronal slices containing CTX and STR were subsequently incubated ≥1.5h with either vehicle Captisol® or 500nM MLi-2 prior to recording. dLight expression is depicted in blue with Stim in dorsolateral STR. **ii)** Representative 2-pulse stimulation with 4s inter-pulse interval with accompanying dLight response. **B** dLight responses were analyzed together (WT veh vs VKI veh vs WT MLi-2 vs VKI MLi-2) and graphed separately. **i)** VKI responses are increased across increasing stimulus intensities compared to WT responses (2-way ANOVA interaction p=0.0003, genotype p=0.0001, Šídák’s multiple comparisons test @50µA *p*=ns, 100-400 µA *p* >0.05). **ii)** MLi-2 treatment does not alter WT dLight responses (Šídák’s multiple comparisons test *p*=ns). **iii)** MLi-2 treatment reduces elevated dLight responses in VKI brain slices (Šídák’s multiple comparisons test @50 µA *p*=0.07, @100-400 µA *p*<0.05). **iv)** Decay of 1^st^ response is not different between VKI and WT brain slices and is not sensitive to MLi-2 treatment (Kruskal-Wallis *p*=0.66). **v)** PPR at 4s interval is not different between VKI and WT brain slices and is not sensitive to MLi-2 treatment (Kruskal-Wallis *p*=0.69).

**Figure 6.**
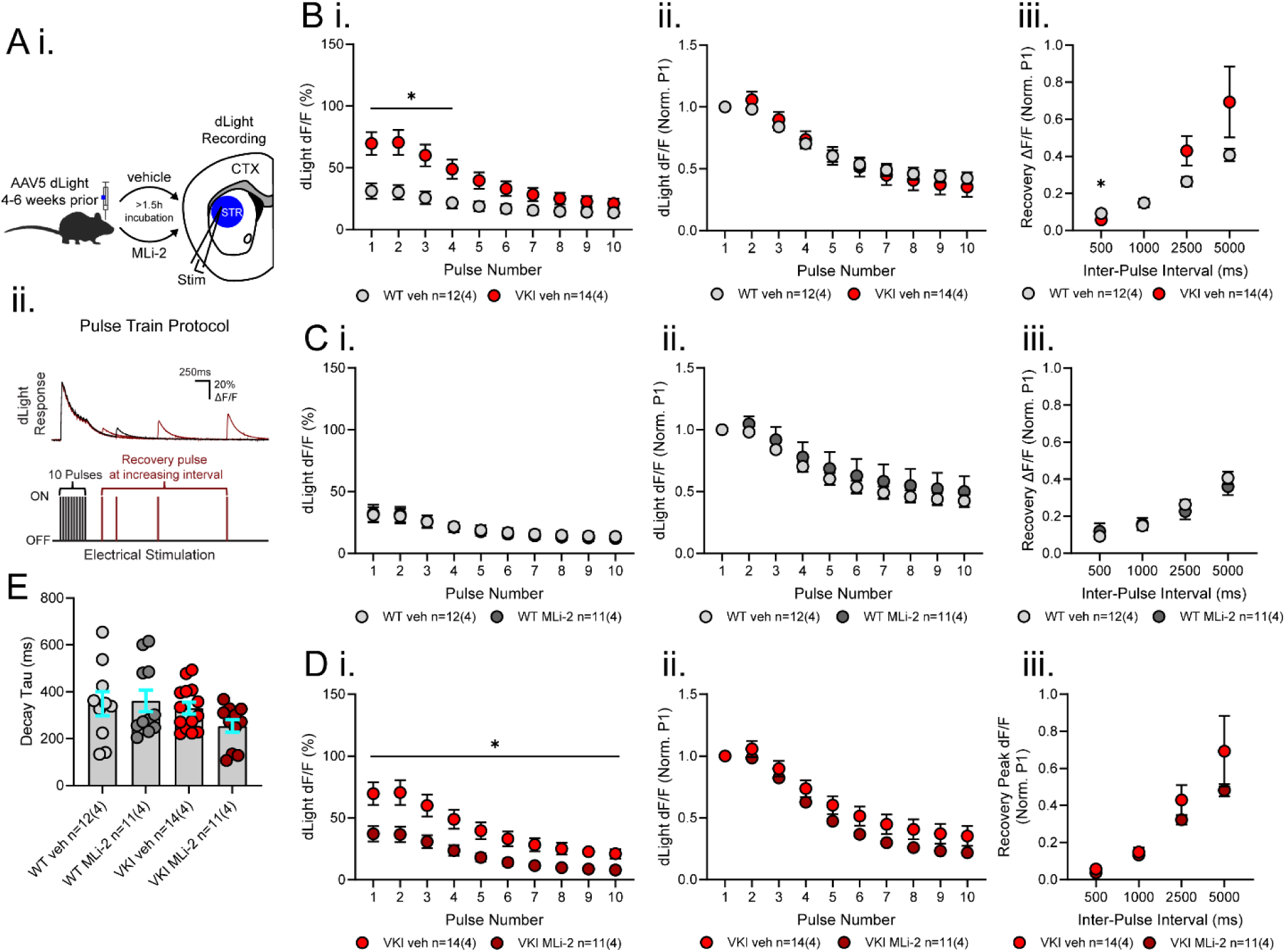
Striatal dopamine release with train stimulation is larger in VKI brain slices and sensitive to acute MLi-2 treatment. **A i)** Graphic depicting stereotaxic injection of AAVs encoding for dLight into the dorsolateral STR 4-6 weeks prior to acute slice preparation. Coronal slices containing CTX and STR were subsequently incubated >1.5h with either vehicle Captisol® or 500nM MLi-2 prior to recording. dLight expression is depicted in blue with Stim in dorsolateral STR. **ii)** Representative response to pulse train stimulation (10x10Hz with repeated 4x with increasing 11^th^ recovery pulse interval). **B-D** dLight response parameters are shown separated by genotype in vehicle treatment and within genotypes by treatment condition. Statistics were performed using 2-way ANOVA on WT veh vs VKI veh vs WT MLi-2 vs VKI MLi-2 with relevant post-hoc comparisons. **B i)** dLight response size is significantly higher in VKI vs WT vehicle-treated brain slices (2-way ANOVA interaction *p*<0.0001, genotype *p*=0.003, Šídák’s multiple comparisons test @P1-P4 *p*<0.05, P5 *p*=0.06, P6-P10 *p*=ns). **ii)** dLight responses, when normalized to the first peak, are similar between VKI and WT vehicle-treated brain slices (2-way ANOVA interaction *p*<0.0001, genotype *p*=0.2393, Šídák’s multiple comparisons test @P1-P10 *p*=ns). **iii)** Recovery capacity following high frequency stimulation is significantly altered in VKI vs WT vehicle-treated brain slices (2-way ANOVA interaction *p*=0.02, genotype *p*=0.19, Šídák’s multiple comparisons test @500ms *p*=0.04, 1000- 5000ms *p*=ns). **C i)** dLight response peaks in WT brain slices are not altered with acute MLi-2 treatment (Šídák’s multiple comparisons test *p*=ns). **ii)** dLight responses normalized to the first peak are not altered with MLi-2 treament (Šídák’s multiple comparisons test *p*=ns). **iii)** Recovery after train stimulation is not altered with MLi-2 in WT brain slices (Šídák’s multiple comparisons test *p*=ns). **D i)** dLight response peaks in VKI brain slices are significantly reduced with MLi-2 treatment (Šídák’s multiple comparisons test @P1-P10 *p*<0.05). **ii)** Normalized dLight responses in VKI brain slices are not altered with MLi-2 treatment (Šídák’s multiple comparisons test *p*=ns). **iii)** Recovery after train stimulation is not altered with MLi-2 treatment in VKI brain slices (Šídák’s multiple comparisons test *p*=ns). **E)** Decay of pulse train response is not significantly different between VKI vs WT brain slices with vehicle or MLi-2 treatment (1-way ANOVA *p*=0.21).

However, recovery following train stimulation was reduced at smaller intervals and increased at larger intervals. WT responses show amplitude reductions between 3 and 6 months, which are absent in VKI responses (Table S2). There is also a significant change to the pattern of normalized responses to pulse train stimulation with age in VKI striata, but not WT. Decay of responses were not significantly different in either WTs or VKIs when comparing 3- to 6-month-old mice. Therefore, the data indicate dopamine release is progressively increased in VKIs, relative to WT, by 6 months old.

### Increased dopamine release in VKI striata is reversed with acute LRRK2 kinase inhibition

We investigated whether the robustly elevated dopamine release in 6-month-old VKI striata could be targeted with acute LRRK2 kinase inhibition. Slices prepared for dLight recordings were incubated either in vehicle Captisol® or in 500nM of LRRK2 kinase inhibitor, MLi-2, for ≥1.5 hours (Figure 5 A). Elevated VKI dLight responses were reversed by LRRK2 inhibition, without altering decay constants, or 4s interval PPRs (Figures 4-6, and Table S1). Although increased amplitudes were similarly reversed in pulse-train experiments, there were no significant effect of MLi-2 on the pattern of responses to stimulation trains. Further, MLi-2 had no effect on WT dLight responses. In conclusion, acute LRRK2 kinase inhibition with MLi-2 reversed the elevated dopamine release in VKI striata at 6 months, without altering dopamine release in WT.

## Discussion

Previously, we found elevated glutamate transmission in 3- to 4-week-old VKI mouse cortical cultures^23^ and elevated striatal dopamine transmission in slices from VKI mice aged 3 months^22,24^. Here, we find corticostriatal glutamatergic transmission onto SPNs is normal at 1 month, but progressively increases to >50% over WT by 6 months. Intriguingly, total striatal glutamate release measured by iGluSnFR was reduced at 3 months and increased to match WT levels by 6 months. In contrast, striatal dopamine release measured by dLight was markedly elevated in VKI striata by 6 months and reversed by acute LRRK2 kinase inhibition.

SPNs of the dorsolateral striatum are the initial point of convergence for control of behavioural action selection, habit formation, and movement initiation^9,12,39^. These functions are all altered in PD, and pathology is clearest first in nigrostriatal axons^1^. Our data suggest SPNs in VKI mice develop changes to passive membrane properties throughout young-adulthood, which could reflect functional and/or anatomical differences. Membrane capacitance is proportional to membrane area, whereas membrane resistance inversely correlates to process diameter (and usually correlates with capacitance). These properties were bidirectionally altered in 6-month-old VKI SPNs, indicative of a larger soma, neurites, and/or processes, and increased leak channel conductance^40,41^.

We observed a subtle increase in amplitude and reduced decay time of spontaneous glutamate events in 3-month-old VKI SPNs, but no changes to optogenetically-evoked corticostriatal AMPAR or NMDAR currents. This suggests any modestly increased response to quantal release in a proportion of synapses is not sufficient to be detected in large currents generated by activity-dependent release. In light of relatively unchanged corticostriatal transmission in SPNs, it was surprising to see that iGluSnFR measures of total glutamate release were markedly reduced in VKI mice at 3 months. This could be explained by an increased number of active synapses, with lower individual release probabilities, or increased postsynaptic responsiveness. Since we previously quantified no change to glutamate synapse number in VKI mice at 3 months^22^, it may be that increased postsynaptic responsiveness exists as an attempt to compensate for one another (or *vice versa*), thereby keeping transmission onto SPNs within a homeostatic range. Alternatively, glutamate release onto SPNs may be unaltered, but reduced at other non-SPN synapses (e.g., excitatory input to cholinergic interneurons^32,34^), inviting future interrogation by cell-type.

By 6 months, clearer increases in SPN spontaneous event amplitude were accompanied by increased frequency (ie. decreased inter-event intervals). At this age, VKI SPN corticostriatal AMPAR- and NMDAR-current amplitudes were also increased ∼50% over WT. More frequent spontaneous activity and higher PPRs, which we also observe in corticostriatal evoked responses, are usually interpreted as increased probability of presynaptic release. Larger AMPAR and NMDAR currents could also be a result of higher postsynaptic receptor numbers. We and others have reported increased GluA1-containing AMPAR receptor expression with VPS35 D620N and LRRK2 G2019S knock-in^23,42^, but increases to both NMDAR and AMPAR expression, here, would have to be equal to explain the lack of difference in NMDAR:AMPAR current ratios here. Thus, the parsimonious explanation is that of increased probability of release, as we reported *in vitro*^23^. Here, direct measures of total striatal presynaptic release with iGluSnFR found no difference between VKI and WT striata, suggesting that corticostriatal transmission onto SPNs must be selectively increased in VKIs. Alternatively, local electrical stimulation may activate the negative tuning of corticostriatal synapses through presynaptic D2 receptors^43^, which would reduce total striatal glutamate release by iGluSnFR imaging, but not affect the direct activation of corticostriatal release with ChR2. Together, the data reveal an emergent increase in cortical glutamate transmission onto VKI SPNs by 6 months, perhaps to compensate for a reduction in total glutamate release observed at 3 months (or *vice versa*).

While not required for diagnosis, changes to glutamate transmission are observed in PD, and likely impact non-motor and motor features of the disease^5,20,44^. Elevated glutamate transmission is seen early in PD-relevant mice carrying knock-in LRRK2 G2019S mutations, and those overexpressing synuclein^19,30,45–47^. Therefore, the progressive dysregulation of glutamate in young adult VKI mice is consistent with other models of PD, and may represent a direct contribution to (or compensation against) pathophysiological progression to degeneration.

Previously, we found dopamine release was elevated in VKI mice at 3 months by fast-scan cyclic voltammetry (FSCV)^22,24^. This was accompanied by slower decay constants, indicative of reduced DAT activity, and a reduction of DAT protein expression^22,24^. Strikingly, increased dopamine release in 3-month-old VKI slices was rescued by chronic 1-week *in vivo* administration of LRRK2 kinase inhibitor, MLi-2^24^. LRRK2 kinase inhibition also normalized the reduced DAT levels, which we concluded was causal to the rescue^24^. Here, we aimed to determine whether the rescue can be observed with shorter-term LRRK2 inhibition (≥1.5h), indicating that dysregulation of dopamine release machinery is LRRK2 kinase-dependent, and not due to systemic effects of LRRK2 kinase inhibition. As FSCV offers low temporal (10Hz) and spatial resolution (limited by the carbon fiber electrode surface area), we employed dLight1.3b to assay dopamine release across the entire dorsolateral striatum at 20x higher temporal resolution.

Considering our published data, we were surprised that dLight recordings showed no difference between VKI and WT dopamine release at 3 months, other than a reduction in the speed of recovery from train stimulation. In contrast, 6-month-old VKI mice displayed a robust elevation in the amplitude of dLight responses, while decay constants were unaltered. The capacity to re-release dopamine after train stimulation was biphasically altered at this age, with shorter time intervals showing a reduced recovery capacity, and longer time intervals showing an increased recovery capacity. Together, this suggests dopamine release is robustly elevated in VKI by 6 months old, and that dopamine axons repolarise, repackage, and re-release dopamine more readily at intervals >1s. Decay constants, and short-term depression (which would be observed as depletion of PPRs during high-frequency trains) are both indicators of DAT function^48–51^. As we saw no difference in these measures, we posit that increased dopamine release is independent of DAT function, as in LRRK2 G2019S knock-in mice^30^.

Differences between our published 3-month VKI voltammetry^22,24^ and dLight data here maybe explained by biological and/or technical differences. Three data sets on mice from the same founding colony now show elevated dopamine release, albeit at slightly different ages^22,24^. While institutional housing conditions may modify phenotype progression, technical differences between slice treatment might also explain the discrepancy. Here, unlike for previous voltammetry data, slices were bathed in NMDG recovery solution to increase slice viability. Recordings were performed in the vehicle for MLi2, Captisol®, a biopolymer that has the potential to increase neurotransmitter release^52–57^, potentially masking differences between genotypes at 3 months. Disparities in the recording modality might also change the interpretation of results at 3 months. Direct comparison of FSCV and dLight shows both techniques report the expected increase in dopamine release duration and decay with nomifensine or cocaine (blocking DAT); however, peak changes are only observed with FSCV^48^. FSCV relies on the area of a carbon fiber electrode onto which dopamine can diffuse, which will increase under conditions of DAT inhibition, and translate to an increase in the peak of FSCV transients^48^. If the spread of extracellular dopamine is increased in VKI at 3 months this would translate to peak changes in voltammetry, but not dLight. Further investigation of the spread dLight fluorescence may reveal whether diffusion is altered at ages preceding clear increases in dLight peaks in VKI mice. Despite these caveats, repeated assessment of dLight across ages in VKI mice reveals increased dopamine release in young-adulthood, which is sensitive to acute LRRK2 kinase inhibition.

Elevated dopamine release in 6-month VKI was returned to WT levels by ≥1.5h slice treatment with LRRK2 inhibitor MLi-2, relative to vehicle-treated slices from the same brain. This occurred without changes to parameters dictated by DAT or D2 receptors (decay rate / pulse train PPR, and 4s PPR^49,58^). Therefore, we conclude LRRK2 kinase hyperactivity and inhibition can alter dopamine release independent of canonical DAT and D2 receptor activity. Both DAT and D2 receptors appear to be VPS35 cargoes, as is the dopamine vesicular monoamine transporter (VMAT)^23,59,60^. It may be that aberrant LRRK2 kinase activity increases vesicular packing of dopamine via VMAT, as VMAT2 levels are elevated in VKI striatal tissue^22,24^. This aligns with our observation that dopamine release is elevated during pulse-trains without significantly altering release probability (P2/P1 PPR, 100ms intervals).

LRRK2 variants convey the highest genetic risk for PD, with several pathogenic variants linked to clinically-typical familial parkinsonism^21^. Brain scans of LRRK2 PD patients show impaired presynaptic dopamine function^61^, but asymptomatic LRRK2 mutation carriers exhibit higher dopamine turnover, indicating increased release^62^. Dopamine release measured by FSCV is also elevated in 3-month-old LRRK2 G2019S knock-in mice^30^, consistent with our results in VKI mice^22,24^. Early increases in dopamine release decline in LRRK2 knock-in mice^30^, and a similar pattern is observed in α-synuclein overexpressing PD mice^63^. Together, evidence from VKI, LRRK2 and synuclein mouse models are consistent with early elevations in dopamine release in humans^62^.

LRRK2 and VPS35 canonically function in endolysosomal regulation and trafficking, and synaptic vesicles (SVs) are highly specialised, neural-specific, endosomes^64^. As with other membranes, the SV cycle is orchestrated by Rabs, several of which are phosphorylated by LRRK2. This includes the canonically synaptic-vesicle associated Rab3&5^65,66^ and classically-endolysosomal Rab10&12, that have recently been shown to be as enriched as Rab3&5 on SVs^67^. Both VPS35 and LRRK2 can alter synaptic vesicle docking and size^68^, although how remains speculative. Conceptually, LRRK2 phosphorylation of Rab3 might be expected to increase SV availability by facilitating biogenesis, trafficking, and priming of vesicles at the active zone. Rab3 tethering to Rab-Interacting Molecule (RIM)^69^ regulates vesicle fusion, without which DA release is blocked^70^. Other than Rab3, the role of Rabs at SVs is somewhat unclear, and effects of LRRK2-Rab phosphorylation almost entirely unknown. Finally, while Rabs have recently taken centre stage, several SV proteins were previously implicated as LRRK2 substrates; including, auxillin, endophilinA, dynamin, & synapsin^47^, and we show here that LRRK2 kinase inhibition can have rapid effects on dopamine release. Further investigation of VPS35 mutant, and LRRK2 inhibitor effects, on dopamine and glutamate vesicle cycles seems justified.

Given the ubiquitous expression of VPS35 and LRRK2, it likely that onset of PD-like dysfunction and degeneration results from the accumulation of low-level cellular insults over years in mice, or decades in humans. Sustained elevations in striatal neurotransmission, or even compensatory action towards it, likely increases cellular stress. This in turn, could lead to the accumulation of excitotoxic damage which may not be effectively cleared in the presence of the VPS35 (or LRRK2) mutation^71^. Increased energy demands and calcium buffering required by increased synaptic activity may also contribute to mitochondrial dysfunction observed in many PD scenarios, including VPS35 patient- derived neurons^72^. These may eventually lead to axon degeneration in more vulnerable neuronal populations, such as nigrostriatal dopamine neurons.

In summary, we present evidence of progressively elevated neurotransmission in young-adult mice with VPS35 D620N mutations, which we propose will become neurotoxic if sustained. However, acute inhibition of LRRK2 kinase rapidly reverses the increased dopamine release, offering hope as a viable neuroprotective strategy. Targeting early neuronal hyperactivity may be the optimal therapeutic window for disease-modifying PD treatments, especially for those known to carry PD-causal gene variants.

## Acknowledgements

**Author Contributions**

A.K. performed most animal surgeries and experiments, contributed to the experimental design and curation, performed data analysis, and co-wrote the manuscript with A.J.M.

C.A.K. contributed intellectually to the interpretation of findings and training of A.K.

N.K. performed initial animal surgeries and contributed to the training of A.K.

S.C. collected a subset of iGluSnFR recordings

E.H. prepared slices and collected recordings for a subset of iGluSnFR experiments

J.C.B. performed animal surgery for a subset of iGluSnFR experiments

M.P.P. contributed to the experimental design and analysis of iGluSnFR experiments and to the training of A.K.

A.J.M. contributed experimental analytical design and curation, training of A.K., and co-wrote the manuscript with A.K.

