## Supplemental Tables and Figures for "Emergent glutamate & dopamine dysfunction in VPS35_(D620N)_ knock-in mice and rapid reversal by LRRK2 inhibition"

**Supplementary Material**

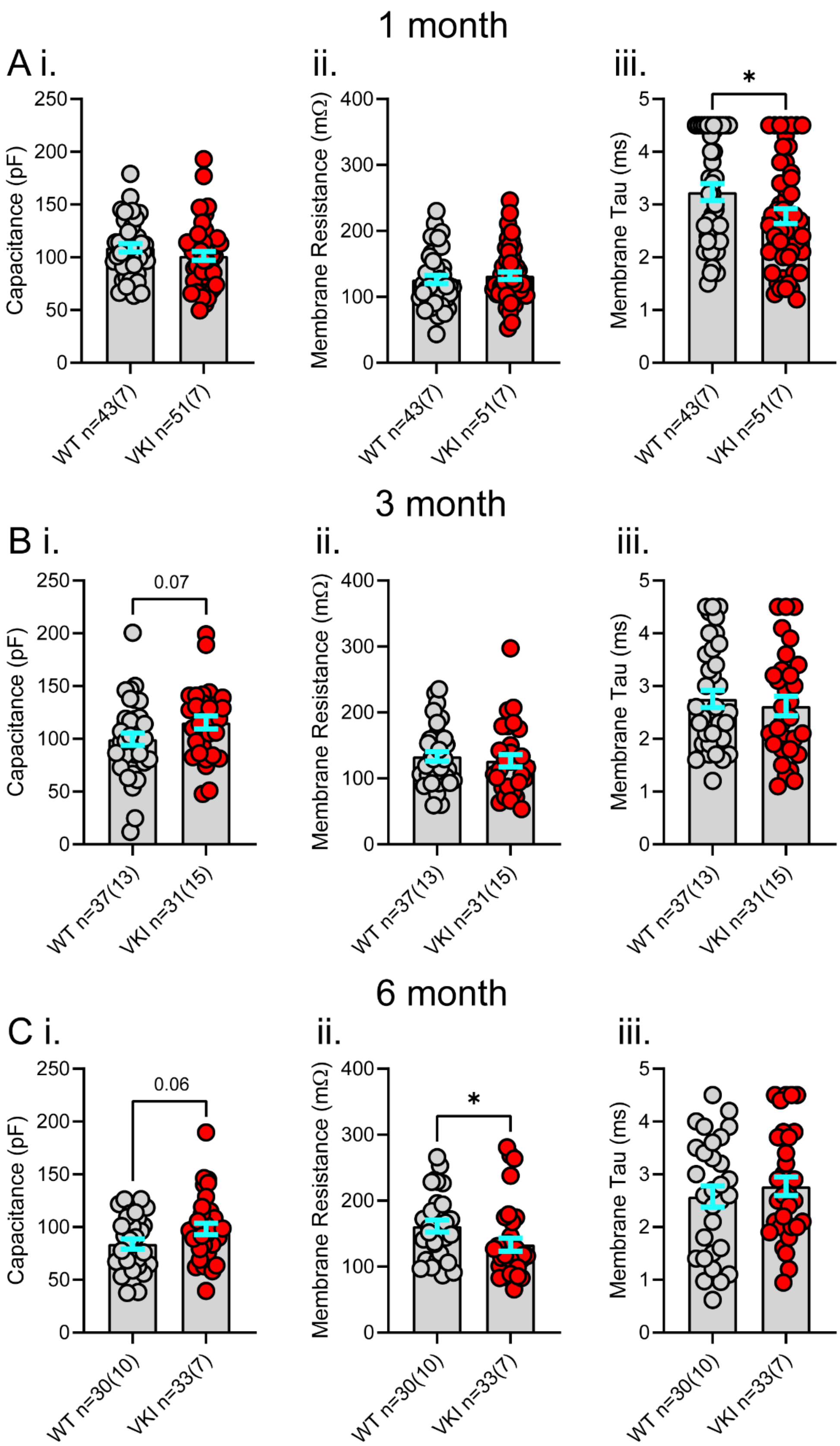

**Figure S1. Membrane properties of dorsolateral striatum spiny projection neurons (SPNs) are altered in VKI mice in an age-dependent manner.**

**Table S1. Additional post-hoc analyses of WT vs VKI measures**

<table>
<tr><th>Measure</th><th>Test Type</th><th>Significance</th><th>Post-Hoc Test Type (if p<0.10)</th><th colspan="3">Post-Hoc Significance (if p<0.10)</th></tr>
<tr><td>1m sEPSC Amplitude</td><td>2-way ANOVA</td><td>Interaction p=0.9996<br>Genotype p=0.2162</td><td>-</td><td colspan="3">-</td></tr>
<tr><td>3m sEPSC Amplitude</td><td>2-way ANOVA</td><td>Interaction p=0.0202<br>Genotype p=0.2050</td><td>Šídák's multiple comparisons test</td><td colspan="3">8-11pA – ns<br>12pA – 0.0378<br>>13 – ns</td></tr>
<tr><td>6m sEPSC Amplitude</td><td>2-way ANOVA</td><td>Interaction p<0.0001<br>Genotype p=0.0188</td><td>Šídák's multiple comparisons test</td><td colspan="3">8-10pA – ns<br>11pA – 0.0710<br>12pA – 0.0005<br>13-15pA – <0.0001<br>16-17pA – <0.001<br>18pA – 0.0157<br>>19pA – ns</td></tr>
<tr><td>1m sEPSC Inter-event Interval</td><td>2-way ANOVA</td><td>Interaction p=0.9933<br>Genotype p=0.9261</td><td>-</td><td colspan="3">-</td></tr>
<tr><td>3m sEPSC Inter-event Interval</td><td>2-way ANOVA</td><td>Interaction p=0.1003<br>Genotype p=0.9492</td><td>Šídák's multiple comparisons test</td><td colspan="3">ns</td></tr>
<tr><td>6m sEPSC Inter-event Interval</td><td>2-way ANOVA</td><td>Interaction p<0.0001<br>Genotype p=0.0565</td><td>Šídák's multiple comparisons test</td><td colspan="3">ns</td></tr>
<tr><td>3m ChR2 Paired-Pulse Ratio</td><td>2-way ANOVA</td><td>Interaction p=0.3415<br>Genotype p=0.2185</td><td>-</td><td colspan="3">-</td></tr>
<tr><td>6m ChR2 Paired-Pulse Ratio</td><td>2-way ANOVA</td><td>Interaction p=0.0011<br>Genotype p=0.0376</td><td>Šídák's multiple comparisons test</td><td colspan="3">P2/P1 – ns<br>P3/P1 – 0.0700<br>P4/P1 – 0.0399</td></tr>
<tr><td>3m iGluSnFR ΔF/F (%)</td><td>2-way ANOVA</td><td>Interaction p<0.0001<br>Genotype p=0.0070</td><td>Šídák's multiple comparisons test</td><td colspan="3">P1 – 0.0767<br>P2 – 0.0591<br>P3 – 0.0739<br>P4 – 0.0762<br>P5 – 0.0784<br>P6 – 0.0764<br>P7 – 0.0865<br>P8 – 0.0850<br>P9 – ns<br>P10 – ns</td></tr>
<tr><td>6m iGluSnFR ΔF/F (%)</td><td>2-way ANOVA</td><td>Interaction p=0.9036<br>Genotype p=0.3989</td><td>-</td><td colspan="3">-</td></tr>
<tr><td>3m iGluSnFR ΔF/F (Norm. P1)</td><td>2-way ANOVA</td><td>Interaction p=0.0063<br>Genotype p=0.9951</td><td>Šídák's multiple comparisons test</td><td colspan="3">ns</td></tr>
<tr><td>6m iGluSnFR ΔF/F (Norm. P1)</td><td>2-way ANOVA</td><td>Interaction p=0.7749<br>Genotype p=0.8681</td><td>-</td><td colspan="3">-</td></tr>
<tr><td>3m iGluSnFR Recovery ΔF/F (Norm. P1)</td><td>2-way ANOVA</td><td>Interaction p=0.8390<br>Genotype p=0.4979</td><td>-</td><td colspan="3">-</td></tr>
<tr><td>6m iGluSnFR Recovery ΔF/F (Norm. P1)</td><td>2-way ANOVA</td><td>Interaction p=0.6251<br>Genotype p=0.1239</td><td>-</td><td colspan="3">-</td></tr>
<tr><td>3m dLight 2-Pulse ΔF/F (%)</td><td>2-way ANOVA</td><td>Interaction p=0.6921<br>Genotype p=0.8337</td><td>-</td><td colspan="3">-</td></tr>
<tr><td>6m dLight 2-Pulse ΔF/F (%)</td><td>2-way ANOVA</td><td>Interaction p=0.0003<br>Genotype p=0.0001</td><td>Šídák's multiple comparisons test</td><td>WT veh vs VKI veh<br>50µA – ns<br>100µA – 0.0043<br>200µA – 0.0058<br>300µA – 0.0177<br>400µA – 0.0192</td><td>WT veh vs WT MLi-2 – ns</td><td>VKI veh vs VKI MLi-2<br>50µA – 0.0740<br>100µA – 0.0261<br>200µA – 0.0192<br>300µA – 0.0192<br>400µA – 0.0141</td></tr>
<tr><td>6m dLight 2-Pulse P1 Decay</td><td>Kruskal-Wallis</td><td>0.6588</td><td>-</td><td>-</td><td>-</td><td>-</td></tr>
<tr><td>6m dLight 2-Pulse Repeated Release Capacity</td><td>1-way ANOVA</td><td>0.6581</td><td>-</td><td>-</td><td>-</td><td>-</td></tr>
<tr><td>3m dLight Pulse Train ΔF/F (%)</td><td>2-way ANOVA</td><td>Interaction p=0.9944<br>Genotype p=0.8450</td><td>-</td><td colspan="3">-</td></tr>
<tr><td>6m dLight Pulse Train ΔF/F (%)</td><td>2-way ANOVA</td><td>Interaction p<0.0001<br>Genotype p=0.0025</td><td>Šídák's multiple comparisons test</td><td>WT veh vs VKI veh –<br>P1 – 0.0108<br>P2 – 0.0113<br>P3 – 0.0148<br>P4- 0.0286<br>P5 – 0.0595<br>P6-P10 – ns</td><td>WT veh vs WT MLi-2 – ns</td><td>VKI veh vs VKI MLi-2 –<br>P1 – 0.0361<br>P2 – 0.0409<br>P3 – 0.0452<br>P4 – 0.0451<br>P5 – 0.0413<br>P6 – 0.0398</td></tr>
</table>

| Measure | Test Type | Significance | Post-Hoc Test Type (if *p*<0.10) | Post-Hoc Significance (if *p*<0.10) | | |
|---|---|---|---|---|---|---|
| | | | | | | P7 – 0.0379<br>P8 – 0.0342<br>P9 – 0.0331<br>P10 – 0.0324 |
| 3m dLight Pulse Train ΔF/F (Norm. P1) | 2-way ANOVA | Interaction *p*=0.6877<br>Genotype *p*=0.4482 | - | - | | |
| 6m dLight Pulse Train ΔF/F (Norm. P1) | 2-way ANOVA | Interaction *p*<0.0001<br>Genotype *p*=0.6198 | Šídák's multiple comparisons test | WT veh vs VKI veh – ns | WT veh vs WT MLi-2 – ns | VKI veh vs VKI MLi-2 – ns |
| 3m dLight Pulse Train Recovery ΔF/F (Norm. P1) | 2-way ANOVA | Interaction *p*=0.8149<br>Genotype *p*=0.0594 | Šídák's multiple comparisons test | 500ms – 0.0297<br>1000-5000ms – ns | | |
| 6m dLight Pulse Train Recovery ΔF/F (Norm. P1) | 2-way ANOVA | Interaction *p*=0.0205<br>Genotype *p*=0.1890 | Šídák's multiple comparisons test | WT veh vs VKI veh – 500ms – 0.0446<br>>1000ms – ns | WT veh vs WT MLi-2 – ns | VKI veh vs VKI MLi-2 – ns |
| 6m dLight Pulse Train Decay Tau | 1-way ANOVA | 0.2052 | - | - | - | - |

**Table S2. Additional analyses of 3m vs 6m effects on WT and VKI measures**

| Measure | Test Type | Significance | Post-Hoc Test Type (if *p*<0.10) | Post-Hoc Comparison | Post-Hoc Significance (if *p*<0.10) |
|---|---|---|---|---|---|
| SPN Membrane Capacitance | Kruskal-Wallis test | 0.0023 | Dunn's multiple comparisons test | 1m WT vs 3m WT | ns |
| | | | | 1m VKI vs 3m VKI | ns |
| | | | | 1m WT vs 6m WT | 0.0115 |
| | | | | 1m VKI vs 6m VKI | ns |
| | | | | 3m WT vs 6m WT | ns |
| | | | | 3m VKI vs 6m VKI | ns |
| SPN Membrane Resistance | Kruskal-Wallis test | 0.0432 | Dunn's multiple comparisons test | 1m WT vs 3m WT | ns |
| | | | | 1m VKI vs 3m VKI | ns |
| | | | | 1m WT vs 6m WT | 0.0616 |
| | | | | 1m VKI vs 6m VKI | ns |
| | | | | 3m WT vs 6m WT | ns |
| | | | | 3m VKI vs 6m VKI | ns |
| SPN Membrane Tau | Kruskal-Wallis test | 0.1099 | - | - | - |
| sEPSC Amplitude | 2-way ANOVA | Interaction *p*<0.0001<br>Genotype/Age *p*<0.0001 | Šídák's multiple comparisons test | 1m WT vs 3m WT | <10pA – ns<br>11-17pA – <0.0001<br>18-19pA – 0.0007<br>20pA – 0.0027<br>21pA – 0.0088<br>22pA- 0.0509<br>>23pA – ns |
| | | | | 1m VKI vs 3m VKI | ns |
| | | | | 1m WT vs 6m WT | 8pA – ns<br>9pA – 0.0109<br>10-21pA - <0.0001<br>22pA – 0.0004<br>23pA – 0.0011<br>24pA – 0.0199<br>>25pA – ns |
| | | | | 1m VKI vs 6m VKI | ns |
| | | | | 3m WT vs 6m WT | ns |
| | | | | 3m VKI vs 6m VKI | ns |
| sEPSC Inter-event Interval | 2-way ANOVA | Interaction *p*<0.0001<br>Genotype/Age *p*<0.0001 | Šídák's multiple comparisons test | 1m WT vs 3m WT | <200pA – ns<br>250pA – 0.0212<br>300-400pA – <0.01<br>450-500pA – <0.001<br>550pA -650pA - <0.01<br>700-850pA - <0.05<br>900pA – 0.0559<br>>950pA - ns |
| | | | | 1m VKI vs 3m VKI | ns |
| | | | | 1m WT vs 6m WT | <100pA – ns<br>150pA – 0.0300<br>200pA – 0.0018<br>250-750pA – <0.0001<br>800-850pA – <0.001<br>900-1050pA – <0.01<br>1100-1200 - <0.05<br>1250-1300 - <0.10<br>>1350pA - ns |
| | | | | 1m VKI vs 6m VKI | ns |
| | | | | 3m WT vs 6m WT | ns |
| | | | | 3m VKI vs 6m VKI | ns |
| sEPSC Decay Tau | Kruskal-Wallis test | 0.0075 | Dunn's multiple comparisons test | 1m WT vs 3m WT | ns |
| | | | | 1m VKI vs 3m VKI | 0.0319 |
| | | | | 1m WT vs 6m WT | ns |
| | | | | 1m VKI vs 6m VKI | ns |
| | | | | 3m WT vs 6m WT | ns |
| | | | | 3m VKI vs 6m VKI | 0.0181 |
| ChR2-PSC Paired-Pulse Ratio | 2-way ANOVA | Interaction *p*<0.0001<br>Genotype/Age *p*=0.0022 | Šídák's multiple comparisons test | 3m WT vs 6m WT | ns |
| | | | | 3m VKI vs 6m VKI | P2/P1 – ns<br>P3/P1 – 0.0005<br>P4/P1 – 0.0002 |

| Measure | Test Type | Significance | Post-Hoc Test Type (if $p$<0.10) | Post-Hoc Comparison | Post-Hoc Significance (if $p$<0.10) |
|---|---|---|---|---|---|
| AMPA Current | 1-way ANOVA | 0.0829 | Tukey's multiple comparisons test | 3m WT vs 6m WT | ns |
| | | | | 3m VKI vs 6m VKI | ns |
| NMDA Current | 1-way ANOVA | 0.0647 | Tukey's multiple comparisons test | 3m WT vs 6m WT | ns |
| | | | | 3m VKI vs 6m VKI | ns |
| NMDA/AMPA Ratio | 1-way ANOVA | 0.0290 | Dunn's multiple comparisons test | 3m WT vs 6m WT | ns |
| | | | | 3m VKI vs 6m VKI | ns |
| AMPA Rectification Index | 1-way ANOVA | 0.2455 | - | - | - |
| iGluSnFR ΔF/F (%) | 2-way ANOVA | Interaction $p$<0.0001<br>Genotype/Age $p$<0.0001 | Šídák's multiple comparisons test | 3m WT vs 6m WT | P1 – 0.0091<br>P2 – 0.0136<br>P3 – 0.0162<br>P4 – 0.0165<br>P5 – 0.0175<br>P6 – 0.0157<br>P7 – 0.0169<br>P8 – 0.0151<br>P9 – 0.0171<br>P10 – 0.0165 |
| | | | | 3m VKI vs 6m VKI | ns |
| iGluSnFR ΔF/F (Norm. P1) | 2-way ANOVA | Interaction $p$=0.1313<br>Genotype/Age $p$=0.8747 | - | - | - |
| iGluSnFR Recovery ΔF/F (Norm. P1) | 2-way ANOVA | Interaction $p$=0.8714<br>Genotype/Age $p$=0.1635 | - | - | - |
| iGluSnFR Decay Tau | Kruskal-Wallis test | 0.0009 | Dunn's multiple comparisons test | 3m WT vs 6m WT | ns |
| | | | | 3m VKI vs 6m VKI | 0.0046 |
| dLight 2-Pulse ΔF/F | 2-way ANOVA | Interaction $p$<0.0001<br>Genotype/Age $p$=0.0006 | Šídák's multiple comparisons test | 3m WT vs 6m WT | 50 μA – 0.0189<br>100μA – 0.0050<br>200μA – 0.0017<br>300μA – 0.0023<br>400μA – 0.0020 |
| | | | | 3m VKI vs 6m VKI | ns |
| dLight 2-Pulse P1 Decay Tau | 1-way ANOVA | 0.6617 | - | - | - |
| dLight 2-Pulse Repeated Release Capacity | 1-way ANOVA | 0.5397 | - | - | - |
| dLight Pulse Train ΔF/F | 2-way ANOVA | Interaction $p$<0.0001<br>Genotype/Age $p$=0.0050 | Šídák's multiple comparisons test | 3m WT vs 6m WT | P1 – 0.0013<br>P2 – 0.0010<br>P3 – 0.0013<br>P4 – 0.0024<br>P5 – 0.0049<br>P6 – 0.0087<br>P7 – 0.0131<br>P8 – 0.0171<br>P9 – 0.0201<br>P10 – 0.0230 |
| | | | | 3m VKI vs 6m VKI | ns |
| dLight Pulse Train ΔF/F (Norm. P1) | 2-way ANOVA | Interaction $p$=0.0011<br>Genotype/Age $p$=0.0961 | Šídák's multiple comparisons test | 3m WT vs 6m WT | P2 – 0.0368<br>P3 – 0.0459<br>P4 – 0.0615<br>P5 – 0.0705<br>P6 – 0.0730<br>P7 – 0.0716<br>P8 – 0.0656<br>P9 – 0.0688<br>P10 – 0.0669 |
| | | | | 3m VKI vs 6m VKI | ns |
| dLight Pulse Train Recovery ΔF/F (Norm. P1) | 2-way ANOVA | Interaction $p$=0.0003<br>Genotype/Age $p$=0.0882 | Šídák's multiple comparisons test | 3m WT vs 6m WT | ns |
| | | | | 3m VKI vs 6m VKI | 500-2500ms – ns<br>5000ms – 0.0133 |
| dLight Pulse Train Decay Tau | 1-way ANOVA | 0.3062 | - | - | - |

### Supplementary Figure Legends

**Figure S1. Membrane properties of dorsolateral striatum spiny projection neurons (SPNs) are altered in VKI mice in an age-dependent manner.**

**A** At 1 month, **i)** VKI SPNs do not show differences in membrane capacitance compared to WT neurons (Mann-Whitney U-test *p*=0.18). **ii)** Membrane resistance is similar between VKI and WT SPNs (Unpaired t-test *p*=0.50), **iii)** Membrane tau is significantly reduced in VKI SPNs vs WT SPNs (Mann-Whitney U-test *p*=0.04). **B** At 3 months, **i)** VKI neurons trend towards significantly higher membrane capacitance measures vs WT neurons (Unpaired t-test *p*=0.07). **ii)** Membrane resistance of neurons is not different between genotypes (Mann-Whitney U-test *p*=0.39). **iii)** Membrane tau is not significantly different between genotypes (Mann-Whitney U-test *p*=0.61). **C** At 6 months, **i)** VKI neurons trend towards significantly higher membrane capacitance measures (Unpaired t-test p=0.06). **ii)** Membrane resistance is significantly lower in VKI vs WT neurons (Mann-Whitney U-test *p*=0.01). **iii)** Membrane tau of neurons is not significantly different between genotypes (Unpaired t-test *p*=0.47). Experimental n = x(y), where x = # of recordings and y = # of mice. Single asterisks denote 0.05>*p*>0.01 and *p* value included where 0.10>*p*>0.05.
